# Can Ebola Virus evolve to be less virulent in humans?

**DOI:** 10.1101/108589

**Authors:** Mircea T. Sofonea, Lafi Aldakak, Luis Fernando Boullosa, Samuel Alizon

## Abstract

This preprint has been reviewed and recommended by Peer Community In Evolutionary Biology (<http://dx.doi.org/10.24072/pci.evolbiol.100022>). Understanding Ebola Virus (EBOV) virulence evolution is not only timely but also raises specific questions because it causes one of the most virulent human infections and it is capable of transmission after the death of its host. Using a compartmental epidemiological model that captures three transmission routes (by regular contact, via dead bodies and by sexual contact), we infer the evolutionary dynamics of case fatality ratio (CFR) on the scale of an outbreak and on the long term. Our major finding is that the virus’s specific life cycle imposes selection for high levels of virulence and that this pattern is robust to parameter variations in biological ranges. In addition to shedding a new light on the ultimate causes of EBOV’s high virulence, these results generate testable predictions and contribute to informing public health policies. In particular, burial management stands out as the most appropriate intervention since it decreases the *R*_0_ of the epidemics, while imposing selection for less virulent strains.

**Impact Summary:** The severe haemorrhagic fever caused by Ebola Virus (EBOV) usually kills more than one infected individual out of two in the absence of treatment, which makes this pathogen one of the most virulent known to humans. The recent outbreak in West Africa (2013-2016) revealed that the virus is able to spread and persist for months across countries. It is often thought that virulence could be due to the fact that the virus is adapted to its reservoir host. Given that microbes evolve rapidly, it is important to determine whether EBOV virulence is likely to decrease as the virus adapts to its human host. To address this problem, we developed a novel mathematical model tailored to EBOV’s life cycle, notably by capturing its three main transmission routes (by regular contact, sexual contact and via dead bodies). We investigated the evolutionary trends of EBOV’s virulence on different time scales (outbreak initiation, short term and long term). Our results reveal that the virulence of EBOV might not be due to the maladaptation of the virus, but could rather originate from its unique life cycle. These results are robust to the parameter values chosen. From a public health perspective, burial management stands out as the main leverage to fight the virulence of EBOV, both on the short and long terms.

## Introduction

Ebola Virus (EBOV) has been a major source of fear since its discovery in 1976. Until 2013, all outbreaks could be traced to spillover from reservoir hosts (Leroy et al., 2005) and were limited in size. This was attributed to EBOV’s extremely high case fatality ratio (CFR), that is the ratio of infected hosts who die from the infection, which we use here as a measure of infection virulence. The dramatic 2013-2016 epidemic in West Africa, which caused more than 28,000 infections and claimed at least 12,000 lives, showed that the virus can persist in the human population for months, therefore raising the question: ‘How will the virulence of Ebola Virus evolve in humans?’ (Kupferschmidt, 2014).

Being an RNA virus, Ebola is prone to rapid evolution (de La Vega et al., 2015) and *in vitro* analyses suggest that the virus has evolved during the outbreak towards an increased tropism for human cells (Urbanowicz et al., 2016). It was first thought that host-parasite interactions should always evolve towards benignity because mild strains seem to have a transmission advantage over strains that kill their hosts rapidly (Méthot, 2012). Since the 1980s, evolutionary biologists have shown that parasite virulence can be adaptive because it may correlate with transmissibility or within-host competitiveness (for a review, see Alizon and Michalakis, 2015). The avirulence theory does remain prevalent in many fields. For instance, some envisage a decrease in EBOV virulence due to host adaptation Kupferschmidt (2014), even though we know the virulence of some human infectious diseases such as HIV or tuberculosis has followed an increasing trend since their emergence (Gagneux, 2012; Herbeck et al., 2012).

Studying virulence as a potentially adaptive trait for the parasite requires encompassing the whole epidemiological life cycle of the infectious agent (Alizon and Michalakis, 2015). In the case of Ebola Virus, most individuals acquire the infection after direct contact with blood, bodily secretions or tissues of other infected humans whether alive or dead (Bausch et al., 2007). This *post-mortem* transmission route cannot be neglected in Ebola Virus epidemics (Chan, 2014), although its magnitude is likely to be low compared to direct transmission (Weitz and Dushoff, 2015). From an evolutionary standpoint, this route might be crucial as well since the timing of life-history events can dramatically affect virulence evolution (Day, 2003). Intuitively, if the virus is still able to transmit after host death, virulence will have a smaller effect on the parasite’s transmission potential. Moreover, there is an increasing evidence that EBOV might also be transmitted through sexual contact even long after the clinical ‘clearance’ of the infection since the virus can persist in the semen for months (Eggo et al., 2015; Thorson et al., 2016; Uyeki et al., 2016).

Will EBOV become more virulent by adapting to humans? To address this question, we use mathematical modelling to determine how case fatality ratio affects the risk of emergence, how it evolves on the long and on the short term. To this end, we introduce an original epidemiological model that captures all three transmission routes of the virus in human populations. We parametrize our model with data from the well-documented 2013-2016 epidemics. We also perform sensitivity analyses on conservative biological ranges of parameter values to verify the robustness of our conclusions.

We find that EBOV undergoes selection for higher case fatality ratios due to its life cycle that includes transmission from hosts after death. This result is robust to most parameter values within biological ranges. We also show that short-term evolutionary dynamics of virulence are more variable but consistently depend on the duration of the incubation period. Finally, we investigate how public health interventions may affect EBOV virulence evolution. We find a direct, but limited, effect of safe burials that may decrease the spread of the virus, while favouring less virulent strains over more virulent ones.

## Methods

For clarity, most of the technical details are shown in Supplementary materials and this section contains verbal description of the model, Figures illustrating the life cycle and a list of parameter values.

### Epidemiological model

Our original compartmental model is based on the classical Susceptible-Exposed-Infected-Recovered (*SEIR*) model, which we enhanced by adding a convalescent class (*C*) that allows for sexual transmission (Abbate et al., 2016) and an infected dead body class (*D*) that allows for *post-mortem* transmission (Legrand et al., 2007; Weitz and Dushoff, 2015). The model is deterministic and does not include additional host heterogeneity, spatial structure or public health interventions.

We incorporated demography through a host inflow term *λ* and a baseline mortality rate *µ*. Susceptible individuals (*S*) can become infected by regular contact with symptomatic infected individuals (*I*) (World Health Organization Ebola Response Team, 2014), by sexual contact with convalescent individuals (*C*) (Mate et al., 2015) and by contact with the dead body of EVD victims, mostly during ritual practices (*D*) (Chan, 2014). The rates at which these events occur are proportional to *β_I_*, *β*_*C*_ and *β*_*D*_ respectively. As in most models (Keeling and Rohani, 2008), we assumed sexual transmission to be frequency-dependent. For non-sexual transmission, we assumed density-dependent transmission following an analysis of the 2013-2016 epidemic (Leander et al., 2016), although performed at a smaller scale than ours. The total population size of live hosts is denoted *N*.

Upon infection, susceptibles move to the so-called ‘exposed’ class (*E*), meaning they are infected but not yet infectious. For Ebola Virus infections, this latency period is also the incubation period, i.e. the time from the contamination to the onset of the symptoms. These symptoms arise at a rate *ω*.

At the end of this latency/incubation period, individual move to the symptomatic infected compartment (*I*). They leave this compartment at a rate *γ*, which we calibrated using the average time elapsed from the onset of the symptoms to death or recovery. We hereafter refer to this as the ‘symptomatic period’. The probability that an infected individual survives the infection is 1 *− α*. The case fatality ratio (CFR), *α*, is our measure of virulence.

We assumed that infected individuals who survive the infection clear the virus from their blood-stream but not from other fluids such as semen and may therefore still transmit EBOV through sexual contacts (Deen et al., 2015). These convalescent individuals (*C*) completely eliminate EBOV at a rate *σ*. Notice that given the severity of the symptoms (Feldmann and Geisbert, 2011) and the fact that the convalescence period is one order of magnitude greater than the symptomatic period, we neglected the sexual transmission from symptomatic infected individuals (*I*).

Based on the current immunological data (Sobarzo et al., 2013), we assumed that full elimination of EBOV from convalescent hosts confers lifelong immunity to recovered individuals (*R*), who do not contribute to the epidemiology.

On the contrary, infected individuals who die from the disease may continue to transmit EBOV if their burial is performed in unsafe conditions, which occurs at a probability *θ*. There is little data from which to estimate this parameter. However, the proportion of EBOV-positive dead bodies has been estimated to decline from 35% to 5% by the end of 2014 in the most populous county of Liberia (Nyenswah et al., 2016). We therefore set the default value *θ* = 0.25. In the analysis, we strongly vary this parameter since it represents an important leverage public health policies have on the epidemics. In the absence of burial team intervention, body fluids from dead bodies remain infectious for a period *ε*^−1^ which is known to be less than 7 days in macaques (Prescott et al., 2015).

Our model, pictured in Figure 1, consists in a set of Ordinary Differential Equations (ODEs) shown in Supplementary Material A.

**Figure 1:**
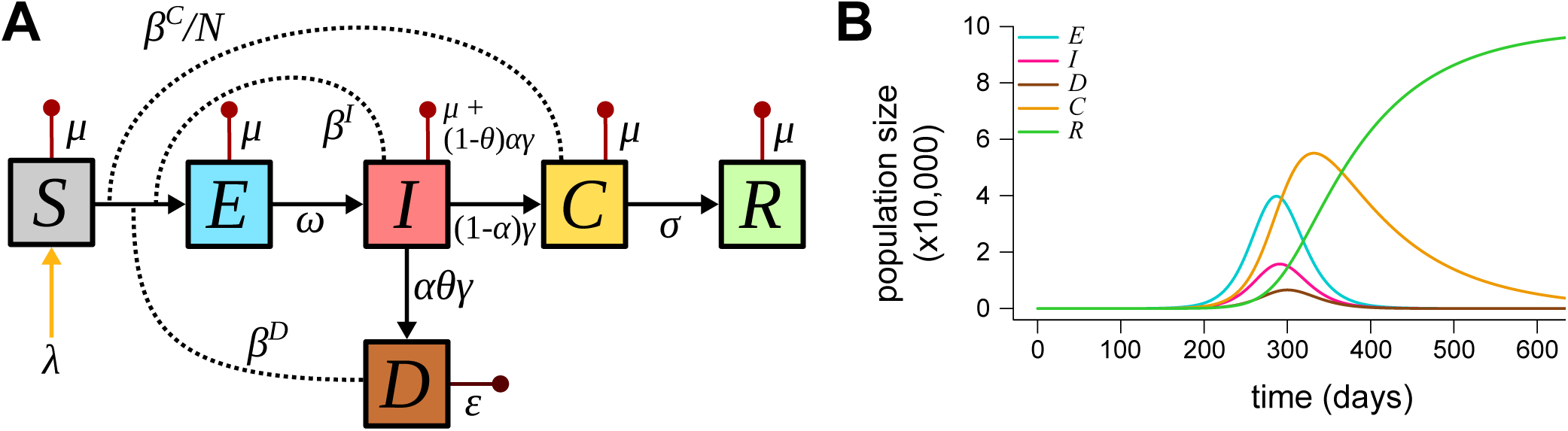
Epidemiology of Ebola Virus in humans. A) Epidemiological life cycle of Ebola Virus in humans and B) Population dynamics for default parameters. *S, E, I, C, R* and *D* are host densities that correspond to the following compartments: susceptible, exposed (infected but not yet infectious), symptomatic infected, convalescent, recovered (immunised) and dead bodies respectively. *N* is the total living population size. Lower-case letters are rate and flow parameters, the description of which is given in Table 1.

Table 1 lists the 11 model parameters. Their values were calibrated using data from the 2013-2016 epidemic in West Africa. We worked at a country level and preferentially chose estimates from the Liberia outbreak, because with approximately 10,000 cumulative cases (World Health Organization, 2016), its magnitude lies in between that of Sierra Leone and Guinea. Demography data from Liberia were obtained from publicly available data from the Central Intelligence Agency (Central Intelligence Agency, 2016). The newborn inflow was set such that the disease free equilibrium matches the country’s population size.

In Supplementary Material C, we calculate the basic reproduction number of EBOV (denoted *R*_0_), which is the average number of secondary infections caused by a single infected individual in a fully susceptible population (Diekmann et al., 1990). By studying the local stability of the system (S1) at the disease free equilibrium, we found that

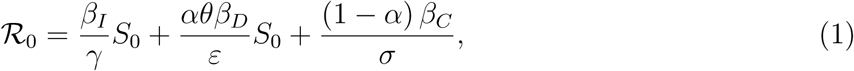

where *S*_0_ = *λ/µ* is the total population size before the epidemic. The three terms in *R*_0_ correspond to each transmission route: from symptomatic individuals (*R*_0,I_ := *β_I_S*_0_*/γ*), infectious bodies (*R*_0,D_ := *αθβ_D_S*_0_*/ε*), and convalescent individuals (*R*_0,C_ := (1 *− α*) *β_C_/σ*). Notice that the incubation period does not affect *R*_0_.

### Transmission-virulence trade-off and estimated values

Trade-offs are a central component of evolutionary model and, without them, predictions tend to be trivial (e.g. viruses should evolve to maximise their transmission rates and minimise their virulence). Although the EBOV life cycle generates constraints that may lead to non-trivial evolutionary outcomes, we do also allow for an explicit trade-off between CFR and transmission rates. Such a relationship has been shown in several host-parasite systems (Alizon and Michalakis, 2015). The case of HIV is particularly well documented (Fraser et al., 2014): viruses causing infections with higher viral loads have increased probability to be transmitted per sexual contact, while causing more virulent (shorter) infections. As a result, there exists an optimal intermediate viral load that balances the virus benefits of transmission with the costs of virulence, thus maximising the number of secondary infections.

In the case of EBOV, there is indirect evidence that viral load is positively correlated with case fatality ratio (CFR) since survivor hosts tend to have lower viral loads than non-survivors (Towner et al., 2004; Crowe et al., 2016). Viral load is thought to correlate with transmission (Osterholm et al., 2015) but demonstrating a clear link is challenging (for HIV, it has required identifying sero-discordant couples in cohorts).

We assumed an increasing relationship between transmission rates and CFR (*α*) such that:

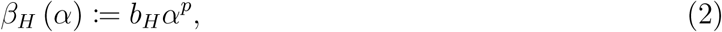

where *b*_*H*_ is a constant factor, *p* ∈ ℝ_+_ is a parameter capturing the concavity of the trade-off curve and *H* stands for one of the compartment *I*, *D* or *C*. The exact value of *p* results from within-host interactions (Alizon and van Baalen, 2005) but one can identify four kinds of trade-offs: *p* > 1 corresponds to an amplifying transmission benefit from increasing CFR, *p* = 1 corresponds to a linear relationship between transmission and CFR, 0 *< p <* 1 corresponds to a saturating transmission benefit from increasing CFR and *p* = 0 is the degenerate case without trade-off. From a biological standpoint, we could expect different transmission routes to have different trade-off shapes (*p*) but, as we show here, our results are largely independent of *p*.

Transmission rates being difficult to estimate (Leander et al., 2016), we indirectly infer the order of magnitudes of *b*_*I*_, *b*_*C*_ and *b*_*D*_ by setting the *R*_0_ close to 2, that is its approximate value for the 2014 epidemic (World Health Organization Ebola Response Team, 2014). Since *R*_0_ is the sum of the epidemiological contributions of each transmission route (see equation (1)), we added the constrain that, according to previous studies, transmission from symptomatic individuals contributes about ten times more than transmission from dead bodies (World Health Organization Ebola Response Team, 2014; Weitz and Dushoff, 2015) and one hundred times more than transmission from convalescent individuals (Abbate et al., 2016). Straightforward calculations (analogous to those done for sensitivity analysis in Supplementary Material E) resulted in attributing the orders of magnitude shown in Table 1.

In the following, the exponent *p* was left undetermined, but its value was set to 0 for the estimation of *b*_*H*_ in the null hypothesis. This leads to *R*_0_ ≈ 1.86, which is very close to the WHO mean estimation for the Liberia epidemic, namely 1.83 (World Health Organization Ebola Response Team, 2014).

### Long term evolution

We used the adaptive dynamics framework (Geritz et al., 1998), which assumes that evolution proceeds by rare and small phenotypical mutations occurring in a monomorphic population that has reached ecological stationarity. Polymorphism is therefore limited to transient dimorphic phases where the ancestor (hereafter called the ‘resident’) and its mutant compete. Depending on the outcome of the competition, the system either goes back to the previous endemic equilibrium or reaches a new monormophic equilibrium. The corresponding dynamical system is shown in Supplementary Material A (system (S2) applied to *n* = 2). Notice that the adaptive dynamics assumptions are particularly consistent with infectious disease dynamics (Dieckmann et al., 2002).

Given the focus of this study, we assumed that the resident and the mutant only differ in their CFR (and in their transmission traits if there was a transmission-virulence trade-off). We denoted the virulence of the mutant and the resident by *α'* and *α* respectively. *α'* was assumed to be close to *α*. Since the mutant is rare by definition, its emergence can be assumed not to affect the resident system. We therefore investigated the local stability of the related endemic equilibrium. This only depended on the local stability of the mutant sub-system (*E*_2_*, I*_2_*, D*_2_*, C*_2_) to which we applied the next-generation theorem (Diekmann et al., 1990; Hurford et al., 2010). This eventually led (see Supplementary Material C.3) to the mutant relative reproduction number

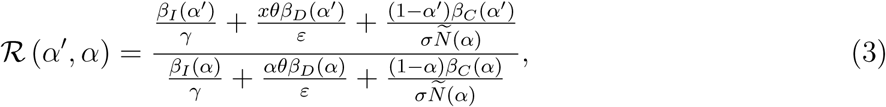

where there total host population size can be approximated (see Supplementary Material B) by

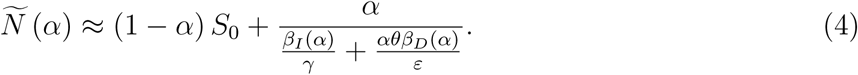

We then calculated the selection gradient by deriving the relative reproduction number (equation (3)) with respect to the mutant’s trait (Otto and Day, 2007). Equating the mutant and resident trait value we eventually found

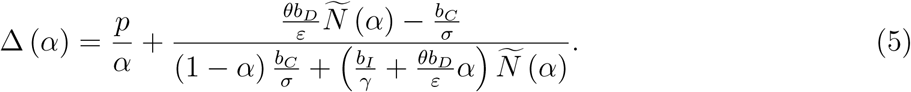

The sign of ∆ indicates the way natural selection acts on the trait depending on the resident’s trait.

### Short term evolution

Viruses evolve so rapidly that evolutionary and epidemiological dynamics may overlap. The epidemiological Price equation framework is designed to predict short-term evolution based on standing genetic variation (Day and Proulx, 2004a; Day and Gandon, 2006).

Practically, we assumed that the parasite population is initially diverse and consists of *n* different genotypes, each genotype *i* being defined by a specific value for several phenotypic traits of interest: the case mortality (*α_i_*), the rate of end of latency (*ω_i_*), the rate of end of the infectious period (*γ_i_*), the rate at which dead bodies cease to be infectious (*ε_i_*), the rate at which convalescent hosts clear the infection (*σ_i_*) and the transmission rates (*β_D,i_*, *β*_*I,i*_ and *β*_*C,i*_).

The dynamics of the populations of interest are described by 4*n* + 1 ODEs that are shown in Supplementary Material A.

After thorough calculations (in Supplementary Material F) we find that, if we neglect mutational bias, mean traits in each compartment vary according to a system of ODEs that involves statistical covariances and variances of traits in the different compartments. The equations involving average CFR are shown in the Results section.

An important assumption of this Price equation approach is that covariance terms are assumed to be constant, which implies that predictions are only valid on the short term. Given that the main limitation of the adaptive dynamics framework relies in its assumption that epidemiological dynamics are fast compared to evolutionary dynamics, combining the two frameworks allows us to get a broader picture.

## Results

### Virulence and emergence

We first consider the risk for an epidemic to occur as a function of EBOV virulence, trade-off shape and burial management. A disease spreads in a fully susceptible population if its reproduction number (*R*_0_) is greater than unity (Anderson and May, 1981). Using our default parameters values (Table 1), we show in Figure 2 that the most virulent EBOV strains are almost always the most likely to emerge, independently of the trade-off exponent *p* and of proportion of unsafe burials *θ*. To have *R*_0_ decrease with *α*, one needs to have neither trade-off nor unsafe burials (*θ* = *p ≈* 0). However, with our default parameter values this decrease is limited.

**Figure 2:**
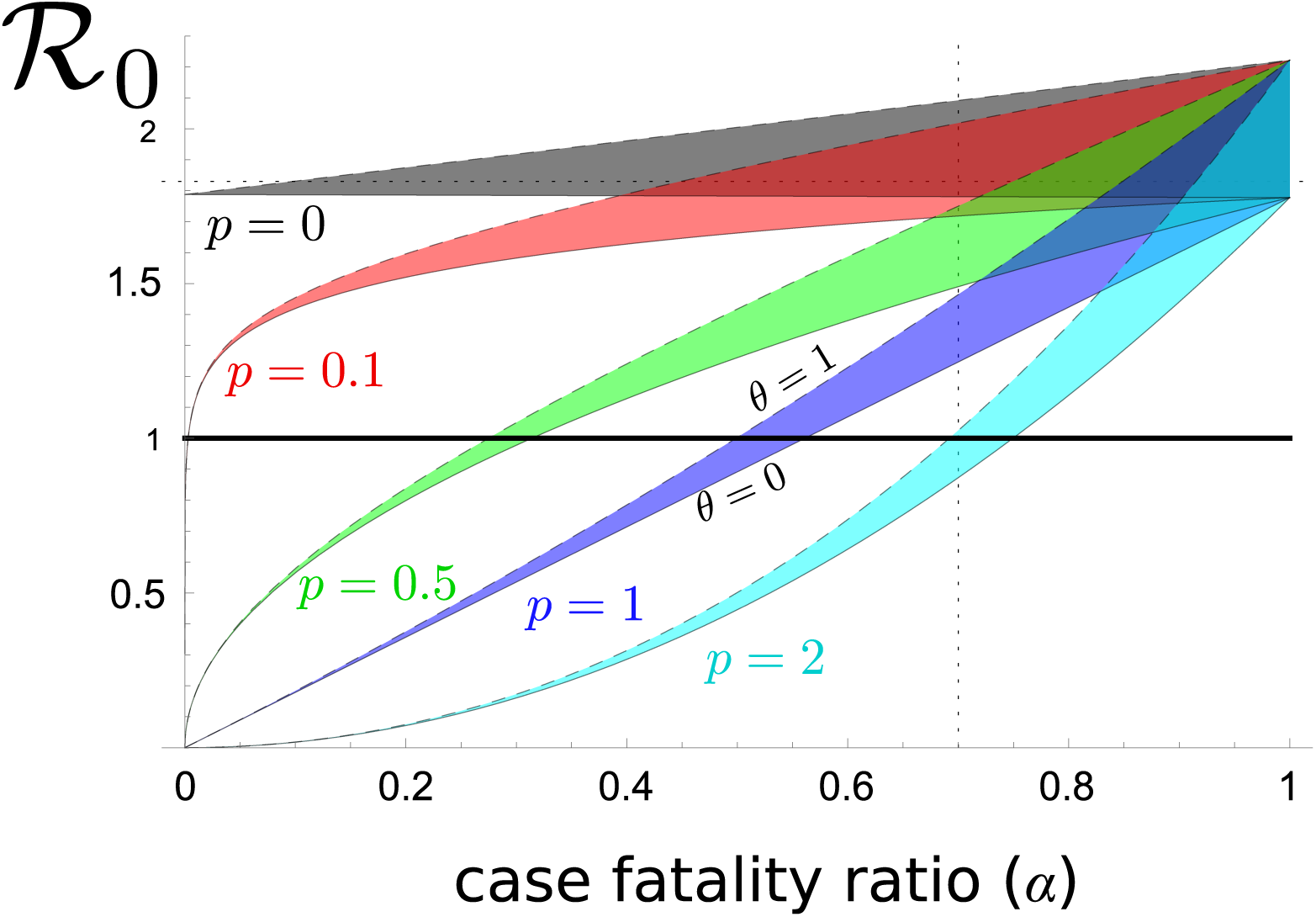
Basic reproduction number as a function case fatality ratio (*α*), unsafe burial ratio and trade-off shape. Colors indicate five trade-off scenarios: absence (*p* = 0, in grey), very weak (*p* = 0.1, in red), concave (*p* = 0.5, in green), linear (*p* = 1, in dark blue), and convex (*p* = 2, in light blue). The width of the coloured regions corresponds to variations in the unsafe burial ratio from completely unsafe (*θ* = 1, dashed upper bound) to completely safe (*θ* = 0, solid lower bound). The intersection between the horizontal line and the colored areas indicates the range of *α*_min_ for each scenario. The dotted gridlines show the *α* and *R*_0_ estimates from the literature. Other parameter values are in Table 1. See Supplementary Material D.4 for more details.

If we focus on the lowest virulence that may lead to an epidemic (denoted *α*_min_), we find that with our default parameter values burial management can prevent emergence (that is bring *R*_0_ below unity by moving vertically in Figure 2) only if the transmission-virulence trade-off is strong enough (the green, blue and cyan curves).

In the following, we will generally assume that EBOV is adapted enough to persist in the human population (*R*_0_ > 1). Since outbreak originates via spillover from reservoir hosts (Leroy et al., 2000), it is likely that the virus is maladapted in the first human infections. However, to capture these dynamics, an evolutionary rescue formalism would be more appropriate given the importance of stochastic events and this is outside the scope of this study (for a review, see Gandon et al., 2013).

### Long-term virulence evolution

If the selection gradient in equation (5) is negative for any CFR (∆(*α*) < 0), then the virus population evolves towards its lowest virulence that allows persistence (that is *α*_min_). If the gradient is always positive (∆(*α*) > 0), the CFR evolves towards 1. Intermediate levels of virulence can only be reached if there exists *α*^*\**^ such that ∆(*α*) *≥* 0 for *α ≤ α^*^*, ∆(*α*) *≤* 0 for *α* ≥ *α*^*\**^ and *R* (*α, α^*^*) < 1 for any *α* ≠ *α^*^*. We show in Supplementary Material D.4 that this occurs only if the proportion of unsafe burials (*θ*) and the trade-off parameter (*p*) are lower than the following boundaries

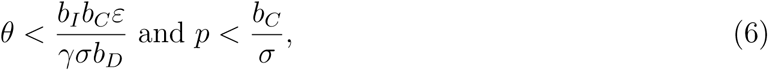

Unless these two conditions are met, the selection gradient is always positive and EBOV is expected to always evolve towards higher case fatality ratios (*α*^*\**^ = 1). Rewriting the first inequality as 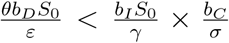 highlights that virulence is favoured by natural selection as soon as the *post mortem* transmission component is greater than the product of the symptomatic and convalescent transmission components.

Figure 3 shows how *α*^*\**^ is numerically affected by a change in burial management (*θ*) and trade-off strength (*p*). Unless the proportion of safe burials is brought below 4%, and unless there is a weak trade-off (*p <* 0.01), CFR will remain high. Intuitively, this double condition can be understood in the following way. If the trade-off is negligible, the CFR is weakly linked to transmission by regular contact and therefore selection on *α* only weakly depends on this component of the life cycle. As a consequence, the value of *θ* governs the relative importance of the two transmission routes that matter. *Post mortem* transmission always favours higher CFR, whereas the sexual transmission route can be maximised for intermediate levels of virulence (see Supplementary Material H.)

**Figure 3:**
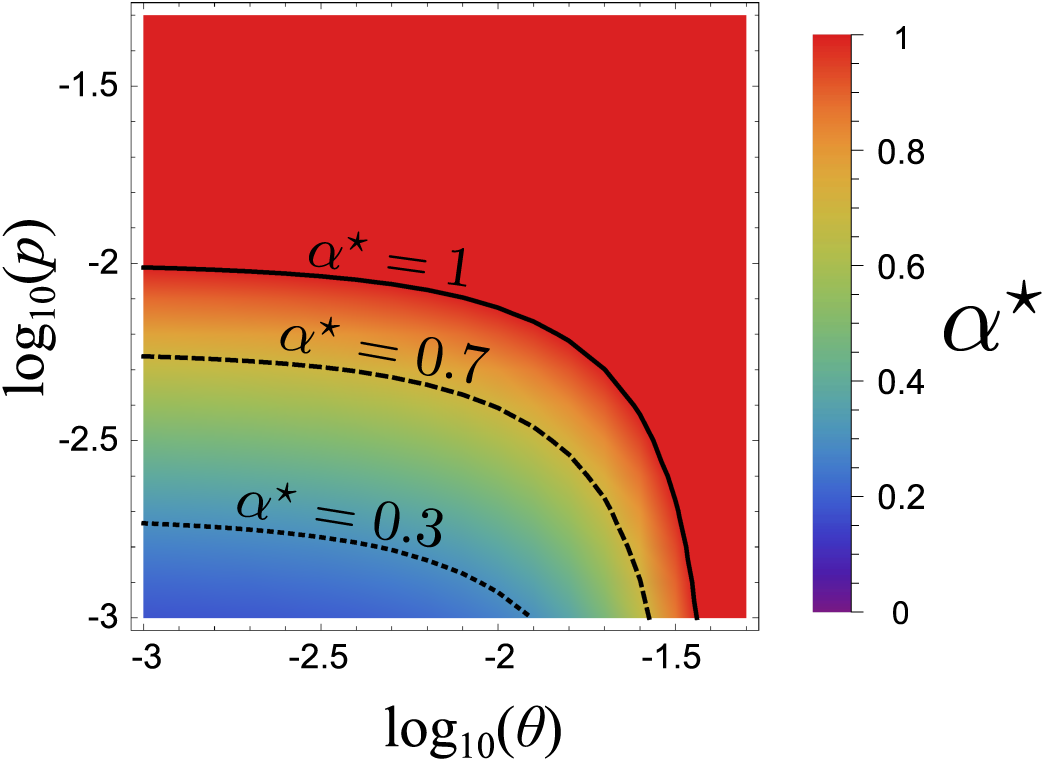
Evolutionary stable virulence (*α^*^*) as a function of unsafe burial ratio (***θ***) and trade-off exponent (*p*). The solid, dashed and dotted lines correspond to *α*^*\**^ = 1, 0.7 and 0.3 respectively. Parameter vlaues are shown in Table 1. See Supplementary Material D.4 for more details.

It was not possible to find an explicit expression for the long-term equilibrium virulence (*α^*^*), but we found it lies in the following interval (Supplementary Material D.5):

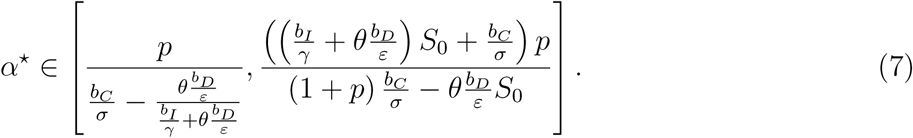

The lower bound of this interval increases with trade-off strength (*p*) and intensity of the *post mortem* transmission route (*θb_D_/ε*). If *post mortem* transmission is strongly reduced, owing to a safer burial management (*θ →* 0), the lower bound simplifies to *pσ/b_C_*. The long-term virulence then appears to be a balance between trade-off strength and sexual transmission intensity, which is consistent with our intuitive explanation of the condition for an intermediate virulence to be selected. In particular, any decrease in the time for convalescent hosts to clear the virus (*i.e.* increase in *σ*) will increase the lower bound for CFR.

To further assess the robustness of these results, we performed a sensitivity analysis by varying the relative importance of each transmission route (regular contact, sexual transmission and transmission from dead bodies), while keeping the total value of *R*_0_ constant. As shown in Figure 4, unless the values of *p* are extremely low, variations in the relative transmission routes is unlikely to be sufficient to move our default value (dotted lines) to the region where low virulences (*e.g. α <* 50%) are favored (green area). To give a quantitative estimate, the relative importance of transmission via sexual contact (on the vertical axis) would need to be about 40 times greater than the current estimates to bring the current estimate above the blue separatrix, which already assumes a low trade-off and a perfect burial management.

**Figure 4:**
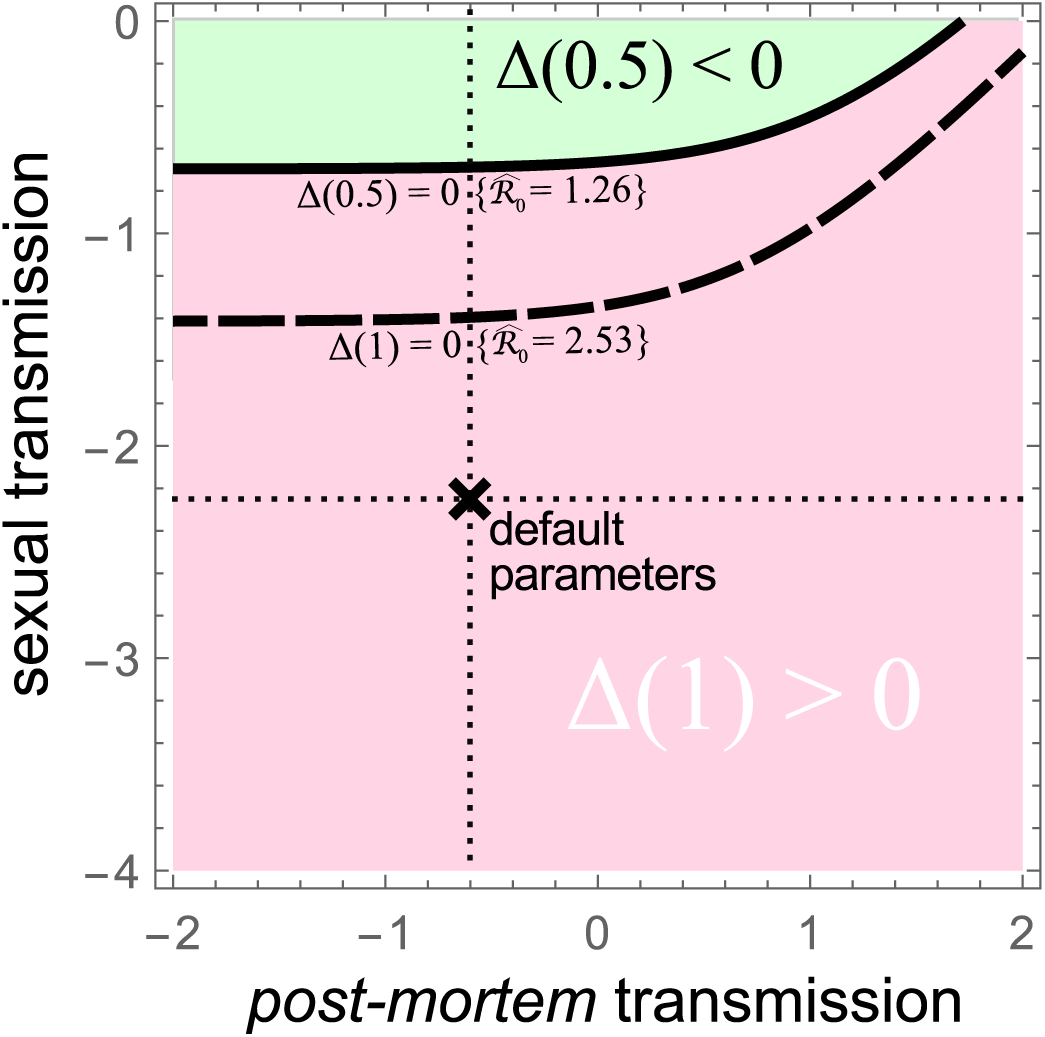
Sensitivity analysis of long-term virulence evolution. The graphic shows the sign of the selection gradient for virulence when varying the relative weight (in orders of magnitude) of the *post-mortem* transmission component (log_10_ (*δ*), on the *x*-axis) and the sexual transmission component (log_10_ (*κ/S*_0_), on the *y*-axis) in the overall transmission of EBOV. When the basic reproduction number is set at its upper bound (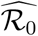 = 2.53, dashed line), the selection gradient at the maximum virulence (*α* = 1) is positive below the dashed line (red area). When the basic reproduction number is set at its lower bound (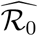 = 1.26, plain line), the selection gradient is also positive for a range of virulence higher than one half (*α* ≥ 50%) in the pink area. It is negative for lower virulences (*α <* 50%) above the dashed line (green area). The unsafe burial proportion and the trade-off exponent are low (*θ* = 0 and *p* = 0.1). See Supplementary Material E for more details.

### Short term evolutionary dynamics

Reaching an evolutionary equilibrium may take time (especially if strains have similar trait values) and the transient dynamics can be non-trivial because the system is non-linear. The Price equation framework provides a way to qualitatively address the initial trends of average trait values, by considering the initial diversity in the virus population.

If we denote by 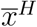 the average value of trait *x* in compartment *H* and by cov *H* (*x, y*) the statistical covariance between traits *x* and *y* in compartment *H* (which becomes the statistical variance var *H* (*x*) if *x ≡ y*), the dynamics of average virulence in the four infected compartments satisfy the following ODEs (see Supplementary Material F for further details):

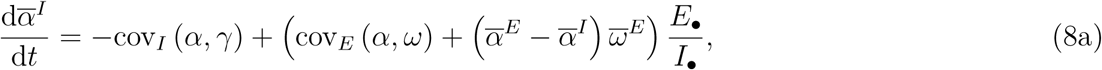

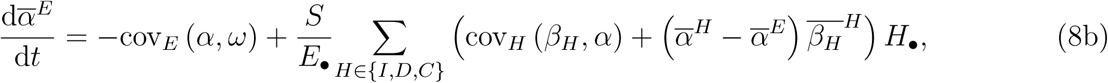

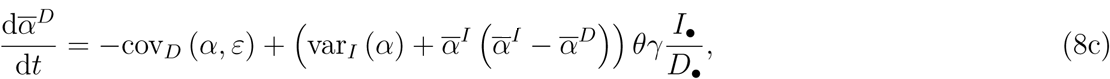

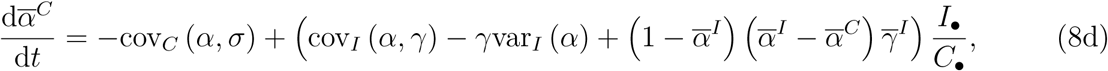

where 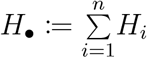 denotes the total size of compartment *H* ≡ *E, I, D, C*.

Focusing on the compartment on which virulence acts, namely the symptomatic individuals, indicates that the short-term evolution of the average virulence 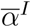 is mainly governed by the correlations between this trait and the symptomatic and latency periods. More explicitly, equation (8a) states that if the most virulent strains induce the longest symptomatic period and/or the shortest latency periods, the average virulence in *I* can be expected to increase at the beginning of the epidemic. Intuitively, newly symptomatic individuals are more likely to have been infected with a highly virulent strain.

Equation (8a) contains a third term proportional to 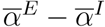, which is more difficult to apprehend. Indeed, 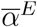 varies as well and follows a complicated ODE that involves not only the correlation with the latency period but also correlations with the transmission rates (equation (8b)). This diversity of components make both 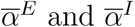 difficult to predict early in the epidemics.

We therefore simulated epidemics numerically according to system (S2). We considered nine scenarios of increasing complexity, three of which are shown in Figure 5 (see Supplementary Material G for more details). During the first six months of an (unmanaged) epidemic, average virulence exhibits wide variations. In most scenarios (panels A and B), it tends to evolve towards the maximum of the range provided by its initial polymorphism. Transient evolution for further virulence in an expanding epidemic have been highlighted by previous models (Day and Proulx, 2004b, e.g.) and studies (Berngruber et al., 2013) but this effect was due to a positive correlation between virulence and transmission rates, which is here not required (Figure 5A). Unsurprisingly, the addition of such correlation amplifies the transient increase in virulence observed during the approximately first 300 days after the onset of the epidemic (Figure 5B).

**Figure 5:**
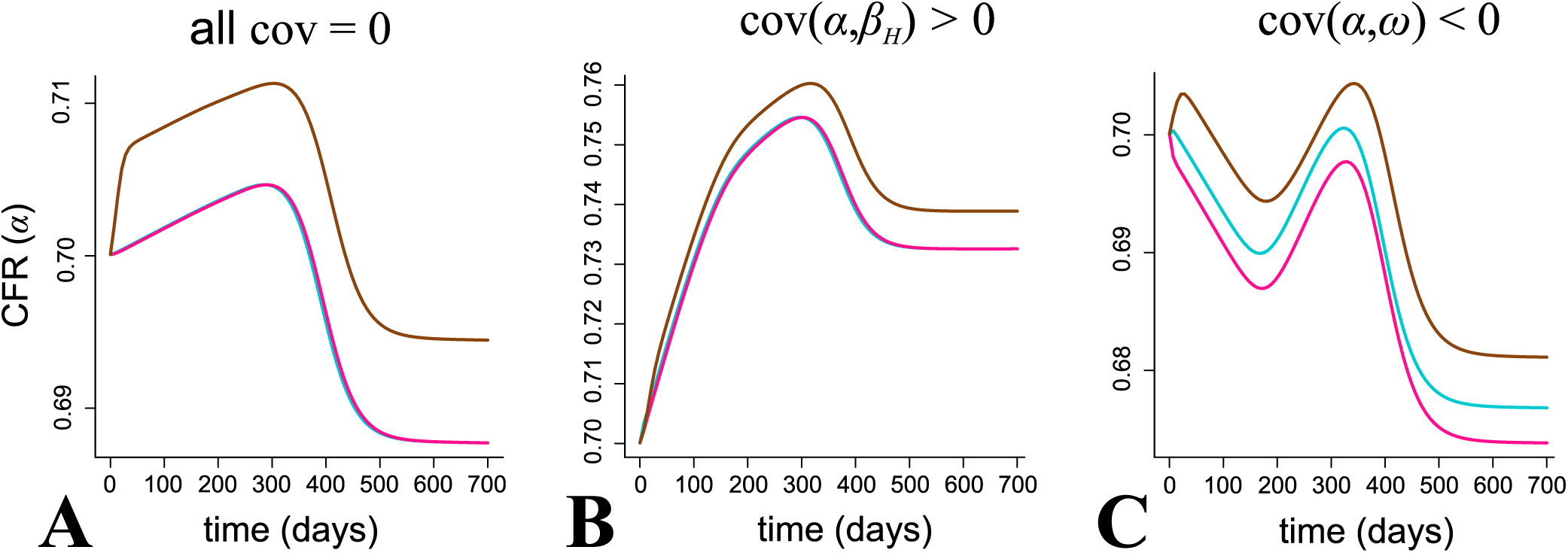
Short-term evolution of CFR with standing genetic variation in three scenarios. A) Without correlations between traits, B) with a positive correlation between CRF and transmission rate and C) with a negative correlation between CRF and latency period. The CFR averaged over the exposed individuals 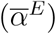 is depicted in cyan, over the symptomatic individuals 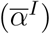 in pink and over the infectious dead bodies 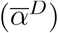 in brown. Parameter values are shown in Table 1. See Supplementary Material G for details about the simulations.

A scenario where average virulence decreases initially is when it is positively correlated with the latency period (Figure 5C). This occurs because less virulent strain have an advantage early in the epidemics by reaching the infectious class earlier. More virulent strains become more frequent again when the value of *D* begins to take off (Figure 5B).

A secondary result shown in these figures is that dead bodies (in brown) always carry more virulent strains on average.

## Discussion

### Virulence could be adaptive for EBOV

Ebola Virus is one of the deadliest human pathogen (Feldmann and Geisbert, 2011). The recent epidemic in West Africa has shown that it can transmit for months months among humans throughout entire countries. As any microbe (especially RNA viruses), it is likely exposed to fast evolution during the course of the epidemic. From a public health standpoint, it is important to predict Ebola virus’ next move and the current hope is that the shift from an emerging to an endemic lifestyle could favour less virulent strains (Kupferschmidt, 2014).

Predicting virulence evolution is usually challenging because we tend to lack details about correlations between epidemiological parameters. Furthermore, even when there is data to estimate a trade-off relationship, its exact shape can have huge quantitative and even qualitative effects on the evolutionary dynamics of the trait (Alizon and van Baalen, 2005; Svennungsen and Kisdi, 2009). Our results stand out because they are robust both to parameter variation in wide biological ranges (World Health Organization Ebola Response Team, 2014, 2015) and also to the type of trade-off assumed (and even to its existence).

In addition to the strong selection on EBOV virulence due to its transmission via dead bodies, another striking result is that decreasing the ratio of unsafe burials is triply effective. First, it decreases the spread of the virus (i.e. its *R*_0_). Second, in the short term, it can help limit a transitory increase in virulence. Indeed, in the first weeks of an epidemic, the sexual transmission route is negligible compared to the other routes that are maximised for the highest CFR values. Third, in the long term, decreasing the proportion of unsafe burials is necessary to shift the selective pressure in favour of less virulent strains.

Overall, EBOV is unlikely to evolve to become less virulent because that would require two conditions. First, the proportion of unsafe burials must be brought to a very low value, which we estimate to be lower than 4%. Second, there must be very little or no genetic relationship between EBOV case fatality ratio and transmission rate. This latter condition is particularly frustrating because it cannot directly be addressed by public health policies. Finally, even if these conditions are met, the level of virulence reached in the long term may still be high, especially if sexual transmission is limited. On a more positive note, results from the Price equation approach show that the virus may experience transitory lower levels of virulence before reaching this maximum via a positive genetic correlation between virulence and incubation period. This is somehow unexpected because this latter parameter does not appear in the calculations related to long-term evolutionary or emergence.

In addition to the strong selection for maximum virulence of EBOV, another striking result is that decreasing the ratio of unsafe burials is triply effective. First, it decreases the spread of the virus (i.e. its *R*_0_). Second, in the short term, it can help limit a transitory increase in virulence. Indeed, in the first weeks of an epidemic, the sexual transmission route is negligible compared to the other routes that are maximised for maximum CFR. Third, in the long term, decreasing the proportion of unsafe burials is necessary to shift the selective pressure in favour of less virulent strains.

### Virulence is a shared trait

In evolutionary biology, virulence is defined as the decrease in host fitness due to the infection (Alizon and Michalakis, 2015). Given the speed at which a pathogen kills its host, EBOV’s virulence can neither be measured as a decrease in instantaneous fecundity (which would be almost zero) nor as an increase of instantaneous mortality rate (which would tend towards infinity or zero depending on the infection outcome). The case fatality ratio (CFR) therefore appears to be the only measurable and epidemiologically relevant proxy for EBOV’s virulence.

We focused on the virus side but, like any infection trait, virulence is also determined by the host and its environment. To predict how virulence will change in the future, we should also consider how hosts may change. In the case of EBOV, it was known before the recent epidemics that some people can become immune to the virus without exhibiting any symptoms (Leroy et al., 2000). The question remains to know if they can also be infectious. More generally, our assumption of life-long protection could be oversimplifying.

Finally, to make predictions on the long term evolution, we need to factor in how the virus population will evolve in response to the variation in the host population’s immune status. Since the immunological status of the host population is determined by that of the virus population, this *de facto* qualifies as a coevolutionary interaction. Earlier models shows that host heterogeneity in resistance and tolerance can lead to a variety of outcomes (Miller et al., 2006; Cousineau and Alizon, 2014). Overall, introducing realistic host heterogeneity, in particular age-dependent or sex-dependent mortality, appears like a relevant extension of this model.

### Spatial structure

Trait evolution is shaped by contact patterns between hosts (Lion and van Baalen, 2008). Regarding the recent Ebola epidemic, the lack of medical personnel and infrastructure in the affected countries played an key role in the spread of the disease as, for example, according to the World Health Organisation, in 2008 Liberia and Sierra Leone had only a density of 0.015 physicians per 1000 inhabitants, when at the same time France had a density of 3.5 and the United States of America 2.4. This was further exacerbated by historical, political and sociological factors (Ali et al., 2016).

It is difficult to predict how explicitly accounting for spatial structure would affect the results. Indeed, it is generally thought that the more ‘viscous’ the population, the more virulence is counter-selected (Boots and Sasaki, 1999). However, the life cycle of the parasite and the host demography can create epidemiological feedbacks that alter this prediction by causing virulence to be maximised for intermediate levels of population structures (Lion and Boots, 2010). This is why predicting virulence evolution in a fully spatially structured model is beyond the scope of this study.

### Testable predictions

One of the underlying assumptions of our model, which could be tested is that the variation we observe in virulence is at least partially controlled by the virus genetics. This could be done by combining virus sequence data with infection traits (virus load or infection outcome) through a phylogenetic comparative approach (Alizon et al., 2010) or a genome wide association study on the virus genome (Power et al., 2017). If virus load is confirmed to be partially controlled by the virus genetics and if, as current evidence suggests, it is correlated with virulence (Towner et al., 2004; Crowe et al., 2016), then studying variations in virus load throughout the 2013-2016 epidemics can help us understand the evolutionary dynamics of virulence. On that note, an experiment consisting in generating pseudovirions based on ancestral or recent EBOV sequences suggests that some of the substitutions observed during the 2013-2016 epidemics may confer increased tropism for human cells (Urbanowicz et al., 2016).

Another result of the short-term evolutionary dynamics analysis is that individuals who contract EBOV from dead bodies should have a higher probability of dying than those infected by contact with living infectious individuals. This could be tested by collecting data from individuals where the transmission route is well documented.

Finally, the remote possibility that lower virulence strains will evolve depends on the existence of a transmission-virulence trade-off. Assessing the shape of this trade-off may be, therefore, very valuable. Note that in the case of EBOV, it is not the exact shape that matters but rather the general trend.

## Conclusion

This evolutionary epidemiology work shows that EBOV’s high virulence, whether it is about emergence, short-term or long-term dynamics, can be explained by its particular life cycle that mixes parasitism and parasitoidism (*post mortem* transmission). Unfortunately, any long term decrease in virulence is unlikely for West African strains at any time scale, although increasing the safe burial proportion appears to be an optimal response in both the short and long terms.

## Authors’ Contributions

Conceived the project: all Performed the analysis: MS Wrote the article: all

**Table 1:** Parameter list, description and default values. See the main text for further details about the calibration of the transmission constants. Note that the sexual transmission constant is higher because it involves frequency-dependent transmission. ind stands for individuals. When used in the main text or in the appendix, these estimated values are denoted by a hat.

| Notation | Description | Default value | Reference or inference |
| --- | --- | --- | --- |
| $\lambda$ | Host inflow | $2 \cdot 10^2 \text{ ind.day}^{-1}$ | such that $\lambda/\mu \approx 4 \cdot 10^6$, (Central Intelligence Agency, 2016) |
| $\mu$ | Host baseline mortality rate | $4.5 \cdot 10^{-5} \text{ day}^{-1}$ | (Central Intelligence Agency, 2016) |
| $b_I$ | Regular contact transmission (from infectious hosts) factor | $10^{-7} \text{ ind}^{-1}.\text{day}^{-1}$ | with the constrain $\mathcal{R}_{0,I} \approx 10 \mathcal{R}_{0,D}$, $\mathcal{R}_{0,I} \approx 10^2 \mathcal{R}_{0,C}$ and $\mathcal{R}_0 \approx 1.8$ (World Health Organization Ebola Response Team, 2014; Weitz and Dushoff, 2015; Abbate et al., 2016) |
| $b_C$ | Sexual transmission (from convalescent hosts) factor | $10^{-4} \text{ day}^{-1}$ | |
| $b_D$ | *Post mortem* transmission (from dead hosts) factor | $10^{-8} \text{ ind}^{-1}.\text{day}^{-1}$ | |
| $\omega$ | Inverse of latency period | $10^{-1} \text{ day}^{-1}$ | (World Health Organization Ebola Response Team, 2014) |
| $\alpha$ | Case fatality ratio | 0.7 | (World Health Organization Ebola Response Team, 2014) |
| $\gamma$ | Inverse of symptomatic infectious period | $0.25 \text{ day}^{-1}$ | (Abbate et al., 2016) |
| $\theta$ | Unsafe burial proportion | 0.25 | (Nyenswah et al., 2016) |
| $\varepsilon$ | Inverse of *post mortem* infectious period | $10^{-1} \text{ day}^{-1}$ | (Prescott et al., 2015) |
| $\sigma$ | Elimination rate of convalescent hosts | $10^{-2} \text{ day}^{-1}$ | (Uyeki et al., 2016; Abbate et al., 2016) |

